# Interaction of micropatterned topographical and biochemical cues to direct neurite growth from spiral ganglion neurons

**DOI:** 10.1101/2021.03.10.434606

**Authors:** Kristy Truong, Braden Leigh, Joseph T. Vecchi, Reid Bartholomew, Linjing Xu, C. Allan Guymon, Marlan R. Hansen

## Abstract

Functional outcomes with neural prosthetic devices, such as cochlear implants, are limited in part due to physical separation between the stimulating elements and the neurons they stimulate. One strategy to close this gap aims to precisely guide neurite regeneration to position the neurites in closer proximity to electrode arrays. Here, we explore the ability of micropatterned biochemical and topographic guidance cues, singly and in combination, to direct the growth of spiral ganglion neuron (SGN) neurites, the neurons targeted by cochlear implants. Photopolymerization of methacrylate monomers was used to form unidirectional topographical features of ridges and grooves in addition to multidirectional patterns with 90° angle turns. Microcontact printing was also used to create similar uni- and multi-directional patterns of peptides on polymer surfaces. Biochemical cues included peptides that facilitate (laminin, LN) or repel (EphA4-Fc) neurite growth. On flat surfaces, SGN neurites preferentially grew on LN-coated stripes and avoided EphA4-Fc-coated stripes. LN or EphA4-Fc was selectively adsorbed onto the ridges or grooves to test the neurite response to a combination of topographical and biochemical cues. Coating the ridges with EphA4-Fc and grooves with LN lead to enhanced SGN alignment to topographical patterns. Conversely, EphA4-Fc coating on the grooves or LN coating on the ridges tended to disrupt alignment to topographical patterns. SGN neurites respond to combinations of topographical and biochemical cues and surface patterning that leverages both cues enhance guided neurite growth.

## 1. Introduction

Although implantable devices such as pacemakers, nerve and deep brain stimulators, and cochlear implants (CIs) have been used for decades to restore vital functions, poor integration with target tissue limits overall performance of neural prostheses. For example, hearing provided by CIs is limited by the significant distance between the electrodes of the CI and the spiral ganglion neurons (SGNs) that they stimulate (Leach et al., 2010; Nadol et al., 2001; O’Leary et al., 2009; Pfingst et al., 2011). Thus, the spatial and temporal resolution inherent in the auditory system is poorly simulated by CIs. As such, CIs have inadequate performance with complex auditory tasks such as perceiving voice inflection and word recognition in noisy environments (Rubinstein, 2004; Shannon et al., 2004).

Directed auditory nerve regeneration, assisted by tissue engineering, stands to dramatically increase the fidelity provided by stimulating electrodes. Regeneration of SGN neurites would help overcome both the high current requirements and broad indiscriminate stimulation that hinder CI performance by placing the primary neural receptors in closer proximity to stimulating elements (Leach et al., 2010; O’Leary et al., 2009; Pettingill et al., 2007; Pfingst et al., 2011; Roehm & Hansen, 2005). To be useful, SGN neurite regeneration must be precisely guided to recapitulate the tonotopic organization of the cochlea. Such SGN regrowth will also be necessary to reinnervate regenerated hair cells, as that becomes feasible.

Strategies to direct neurite guidance have commonly relied on the use of patterned bioactive molecules (Branch et al., 1998; Branch, Wheeler et al., 2001; Clarke et al., 2011; Gustavsson et al., 2007; Kijenska-Gawronska et al., 2019; Millet et al.; Tuft et al., 2014). Using techniques such as microcontact printing, surface modification for selective adsorption, and microfluidics, patterns of peptides that guide neurite growth have been created and used as platforms for directed neurite regeneration (Belisle et al., 2008; Fricke et al., 2011; Honegger et al., 2016; Joo et al., 2015; Klein et al.; Wittig et al., 2005). For example, SGN neurites preferentially grow on micropatterned lines of laminin (LN) and other extracellular matrix proteins (Evans et al., 2007). Other studies have used concentration gradients of neurotrophin-3 (NT-3) generated by a microfluidic compartment to guide SGN neurite growth (Wittig et al., 2005). Neurite outgrowth can also be controlled by patterning proteins that repel neurite growth. For example, EphA4-Fc is a chimeric peptide consisting of the chemorepulsive extracellular domain of EphA4 fused to the Fc portion of immunoglobulin that has been shown to repel SGN neurite growth (Brors et al., 2003; Defourny et al., 2013; Tuft et al., 2018).

In addition to micropatterning of bioactive molecules, tissue engineers have used topographical patterning to guide axon regeneration (Johansson et al., 2006; W. Li et al., 2015; Tuft et al., 2013). Recently we demonstrated that micropatterned methacrylate (MA) polymers consisting of parallel lines of ridges and grooves provide robust guidance cues for SGNs and other neurons (Clarke et al., 2011; Tuft et al., 2013). SGN neurite guidance by these topographical patterns depends on feature periodicity (the distance between consecutive ridges or grooves) and amplitude (the depth between a ridge and a groove). Features of high amplitude (e.g. 8 μm) or low periodicity (e.g. 10 μm) provide robust guidance while features of lower amplitude (1 μm) or greater periodicity (e.g. 100 μm) are less effective at guiding SGN neurite growth (Tuft et al., 2013). Additionally, our lab has investigated using multidirectional patterns with continuous 90 degree turns (Tuft et al., 2014). In both cases, the topographical features have been shown to influence, but not completely dictate, SGN neurite pathfinding thereby permitting exploration of synergistic interaction with biochemical cues. These MA polymers are particularly suited for application as nerve regeneration platforms due to their well-documented biocompatibility in neural and other tissues, (Apple & Sims, 1996; Kenny & Buggy, 2003; Tuft et al., 2013; Uludag et al. 2000) their stability, and the precise control of surface topographical features achieved via micro- and nanoscale patterning techniques.

Understanding the chemical and topographical cues that direct neurite growth will provide critical information for designing neural prosthetics materials. Since neural cells respond to a complex milieu of directing mechanisms, developing a fundamental understanding of cell-material interactions that influence nerve growth will be important for development of next generation devices. Here, we explore the extent to which combinations of micropatterned peptide and topographical features interact to direct SGN neurite growth. To do so, we developed techniques to selectively adsorb peptides that guide SGN neurites to the grooves, ridges, or both grooves and ridges of micropatterned MA polymer surfaces. These combined biochemical-topographical substrates allow for the investigation of the relative impact and synergistic effects of these two different types of guidance cues on directed neurite outgrowth. These are among the first studies to explore the response of cells from the spiral ganglion to topographic features and patterning of biochemical guidance cues. This is also unique as we examined how biochemical, cellular, and topographical features interact independently and in combination to affect neurite growth and pathfinding. In total, this work has direct relevance for efforts to precisely guide SGN neurite regeneration.

## 2. Materials and Methods

### 2.1 Glass Slide Pretreatment

Standard 2.54 cm × 7.62 cm glass slides were exposed to oxygen plasma for 3 minutes at 30 W RF power (PDC-001 Harrick Plasma Expanded Cleaner, Ithaca, NY). Directly after exposure to plasma, the slides were immersed in a 1% v/v solution of 3- (trimethoxysilyl)propyl methacrylate (Aldrich) and hexanes (Aldrich) for > 12 hours at room temperature. After removal from the solution the samples were washed with fresh hexanes and dried before being placed in a sealed container, where they were kept until immediately before use.

### 2.2 Generation of micropatterned MA polymers

Micropatterned polymer surfaces were generated using the inherent temporal and spatial control afforded by photopolymerization. Monomer formulations of 40 wt. % hexyl methacrylate (HMA), 59 wt. % hexanediol dimethacrylate (HDDMA), and 1 wt. % 2,2- Dimethoxy-2-phenylacetophenone (DMPA) were used for each polymer sample. A solution of 20 μL was pipetted onto glass functionalized with a silane coupling agent and dispersed evenly by placing on top a 2.54 cm × 2.54 cm square of either a glass-chrome Ronchi ruled photomask (Applied Image Inc.), 90-degree angled photomask (Applied Image Inc.), or plain glass. Samples were then cured with 365 nm light at an intensity of 16 mW/cm^2^ using a high-pressure mercury vapor arc lamp (Omnicure S1500, Lumen Dynamics, Ontario, Canada) for 120 seconds. Parallel Ronchi-rule or analogous repeating 90° angled patterns consisting of alternating transparent and reflective 25 μm wide bands were used as photomasks (Clarke et al., 2011; Tuft et al., 2013; Tuft et al., 2014). By selectively shielding areas underneath the mask from exposure to UV radiation, the rate of the polymerization is modulated locally to generate features on the surface. Raised features or ridges form underneath transparent bands where UV light intensity and the polymerization rate are highest. Due to diffraction and reflection, regions underneath reflective bands are still exposed to UV light, but at much lower intensities. As such, polymer is not generated as quickly under shadowed regions leading to depressed features or grooves. Photomask band spacing specifies feature periodicity while the amplitude of the features is controlled by the timing and intensity of irradiation (Tuft et al., 2013). After polymerization, the photomask is removed and unreacted monomer is rinsed from the channels with ethanol to reveal a micropatterned surface with gradually sloping transitions between ridges and grooves.

Our labs recently demonstrated that these patterned MA substrates enable attachment and survival of a variety of neurons and glial cells, including SGNs and SG Schwann cells, while also directing their growth (Clarke et al., 2011; Tuft et al., 2013). Neurite guidance by micropatterned MA substrates depends on feature amplitude and periodicity (Tuft et al., 2013); more frequent or higher amplitude features induce greater alignment while less frequent or lower amplitude features provide weaker guidance. To provide SGN neurites with a topographic guidance cue sufficient to induce alignment of neurites, but still allow for enhancement or disruption of the alignment by surface peptides, micropatterns with an amplitude of 2 μm and periodicity of 50 μm were chosen.

### 2.3 Micropatterning of proteins

LN was used as a chemoattractive cue and EphA4-Fc as a chemorepulsive cue (Tuft et al., 2018). We used the same micropatterned MA platforms used in neurite guidance studies as stamps to print LN or EphA4-Fc peptides on MA substrates lacking topographic patterns (Fig. 1) (Tuft et al., 2013). Micropatterned MA stamps were immersed in a solution of either LN (20 μg/ml, Sigma-Aldrich Cat#: L2020, Lot number: 121M4043V) or EphA4-Fc (10 μg/ml, R&D Systems). The peptide-‘inked’ stamps were pressed onto a smooth polymer. This method allowed us to establish precise patterns of peptides. We also formed peptide printed topographical materials by applying proteins to a stamp with an unpatterned surface and then printing them onto the ridges of a micropattern, thereby producing substrates with both topographic and biochemical micropatterns (Fig. 1). To assess the contribution of LN working synergistically with groove features, we formed materials with LN adsorbed only in the grooves by placing sufficient LN solution to fill grooves while leaving the ridges without any LN solution. Any excess protein solution was absorbed from the ridges by placing absorbent paper and a glass coverslip on the ridges, leaving the LN solution only in the substrate grooves.

**Figure 1.**
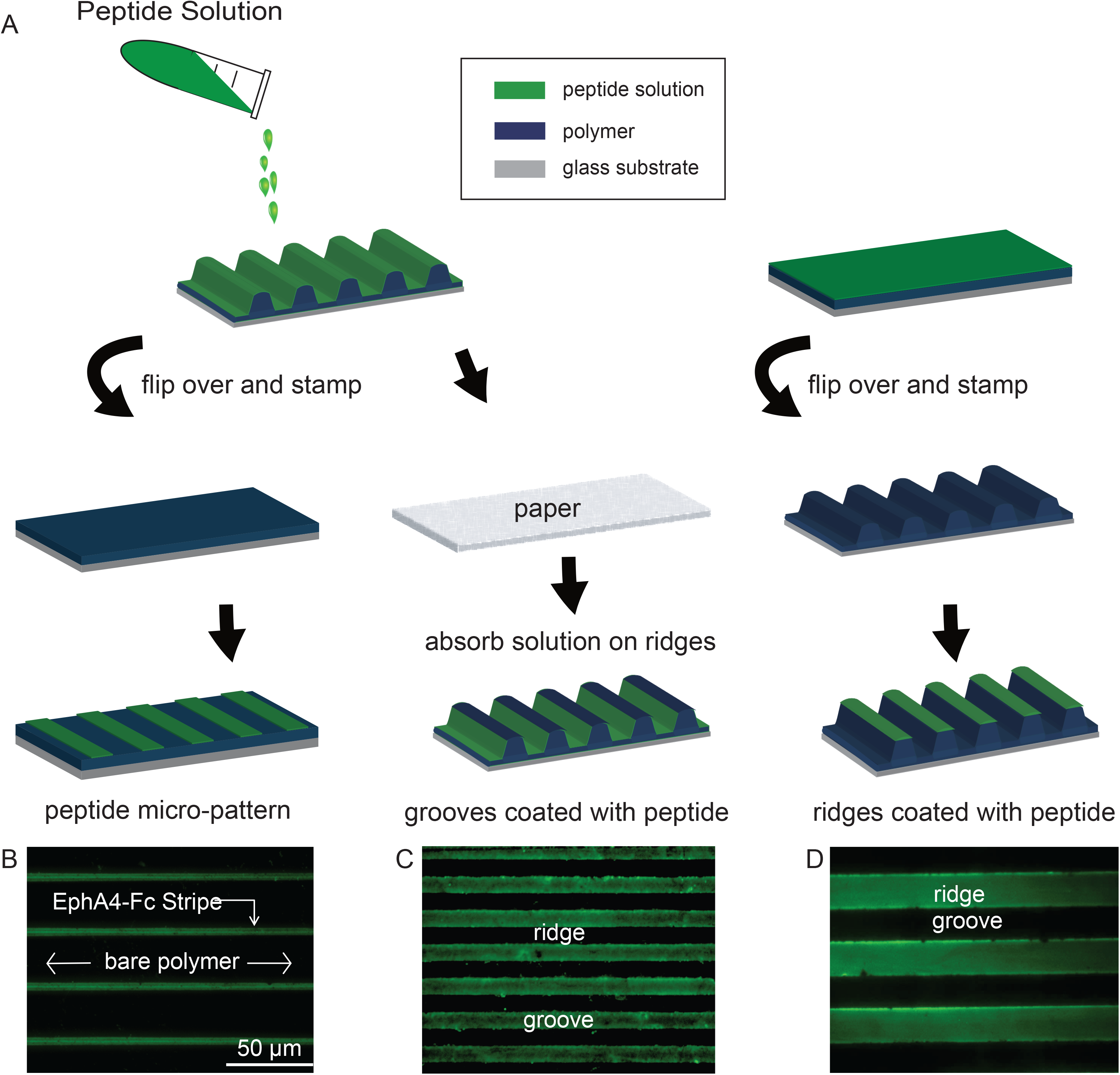
Microcontact printing of peptide patterns. **A**. Schematic drawing representing techniques for selective patterning of peptides on smooth and topographically patterned polymer substrates. **B**. Example of EphA4-Fc pattern printed as stripes on a smooth unpatterned HMA/HDDMA polymer and labeled with Alexa 488 (green) conjugated anti-Fc secondary antibody. **C**. Example of laminin printed into the grooves of a topographically patterned polymer detected by immunolabeling with anti-laminin primary antibody and Alexa 488 conjugated secondary antibody. **D**. Example of EphA-4-Fc printing onto the ridges of a topographically patterned polymer detected by labeling with Alexa 488 conjugated anti-Fc antibody.

### 2.4 SGN cultures and immunofluorescent labeling

The institutional animal care and use committee at the University of Iowa approved all protocols and experimental plans used in this study. Dissociated spiral ganglion cultures protocols were prepared as previously described (Clarke et al., 2011; Hansen et al. 2001; Jeon et al. 2011; Tuft et al., 2013). Briefly cochlear tissues were harvested from postnatal day 4-6 rat pups after rapid decapitation under cold anesthesia. Phosphate buffered saline (PBS) was used as the dissecting buffer. Cultures were plated on various combinations of topographically and/or biochemically patterned substrates (Clarke et al., 2011; Hansen et al., 2001; Jeon et al., 2011; Tuft et al., 2013). Cultures were maintained in the presence of neurotrophin-3 (50ng/ml, Sigma-Aldrich) and brain derived neurotrophic factor (50 ng/ml, R&D Systems) to support neuronal survival (Hansen et al. 2001). After 72 h, the cultures were fixed with 4% paraformaldehyde and immunolabeled with anti-neurofilament 200 (NF200) antibodies (1:400, Sigma-Aldrich, catalog # N0142) followed by an Alexa 488 or Alexa 568 conjugated secondary antibody (1:800, Invitrogen). A subset of slides with microcontact printed peptides were immunolabeled with anti-laminin (1:500, Abcam, catalog # 30320) antibody followed by either Alexa 488 or Alexa 568 conjugated secondary antibody. Another subset of slides was labeled with Alexa 568 conjugated anti-human IgG antibody (1:800, ThermoFisher, A-21090) to detect the Fc portion of the EphA4-Fc chimera peptide.

### 2.5 Determination of SGN neurite behavior to micropatterns

Digital epifluorescence images were captured of the entire polymer surface using the scan slide application of Metamorph software (Molecular Devices) on a Leica DMIRE2 microscope (Leica Microsystem) with a Leica DFC350FX digital camera. The primary outcome measure was neurite alignment to unidirectional and multidirectional biochemical and/or topographic patterns using Image J software (National Institute of Health) as previously described. Secondary outcome measures included: (1) overall neurite length, (2) neurite preference for microfeature (e.g. ridge vs groove or LN vs EphA4-FC stripe) determined as previously described (Tuft et al., 2013). Neurites from 50 randomly selected SGNs were scored for each condition. Each condition was repeated at least 3 times.

For 90-degree patterns, neurite alignment was quantitatively compared using a custom algorithm in ImageJ and MATLAB^®^ to determine the alignment angles of 10 μm segments along the entire length of the neurite, as previously described. In brief, neurite paths were sectioned into 10 μm segments and the alignment angle (θ) difference of each segment relative to the horizontal plane was determined. Alignment to the horizontal is evidenced by angles near 0° while alignment to the vertical yields, near 90°. Angles near 45° suggest that the neurite segment follows the alternating 90° angles and random neurite growth results in neurites evenly distributed across the 0° - 90° spectrum (Tuft et al., 2014).

### 2.6 Statistical Analysis

Differences among experimental groups for data that are normally distributed were determined by a Student’s two tailed t-test for two groups or one-way ANOVA with a post-hoc Tukey analysis for multiple group comparison using SigmaStat software (Systat Software, Inc). Statistical significance for nonparametric data was determined by a Kruskal-Wallis ANOVA on ranks followed by a post-hoc Wilcoxon-Mann-Whitney test. Assuming similar error as in our prior experiments, an a priori power analysis indicated a >95% chance to detect a 20% difference in neurite alignment or length by analyzing 50 neurites for each condition.

## 3. Results

### 3.1 Microcontact printed substrates

Microcontact printing was used to generate patterns of peptides adsorbed to the surface of the MA polymers (Fig. 1). We generated both smooth and topographically patterned substrates printed with either LN or EphA4-Fc peptides. LN is an extracellular matrix protein that facilitates SGN neurite growth (Aletsee et al., 2002; Evans et al., 2007; N. Xu et al., 2012). EphA4-Fc is a chimeric peptide composed of the extracellular domain of EphA4 and the Fc portion of IgG immunoglobulin and functions as a chemorepulsive cue to SGN neurites (Brors et al., 2003; Coate et al., 2012; Defourny et al., 2013; Tuft et al., 2018). To generate micropatterns of peptides with similar patterning as the topographical patterns, peptides were adsorbed to a polymer substrate micropatterned with linear ridges and grooves. The coated polymers were then pressed onto a smooth polymer and a portion of the peptide was transferred to the polymer surface. Immunolabeled and epifluorescence microscopy were used to assess the peptide patterning on the polymer surfaces. Uniform linear strips of the peptides were successfully applied to smooth MA polymers (Fig. 1).

Figure 1 schematically illustrates the application of peptides to either the grooves or ridges of topographically patterned MA polymers. To selectively coat ridges with peptides, a smooth polymer film was coated with the peptide solution. The peptide-coated surface was then gently pressed onto the ridges of a topographically patterned polymer allowing the peptide to transfer to the ridges of the patterned surfaces. Selective coating of grooves was accomplished by first coating a topographically patterned polymer film with a very small volume of peptide solution that remained mostly in the grooves. Any excess peptide solution was quickly removed from the ridges with absorbent tissue. As above, immunofluorescence labeling was used to confirm the selective coating of ridge and groove features (Fig. 1). These results demonstrate the successful application of peptide patterns onto both smooth polymers as well as specific features of topographically patterned polymers.

### 3.2 SGN neurites align strongly to surfaces with LN or topographical micropatterns

We next studied the effect of specific combinatorial peptide and topographic patterns on SGN neurite guidance. The topographical patterns had a periodicity of 50 μm with amplitudes of 2 μm. Based on our prior experience, these topographical patterns are expected to induce modest neurite alignment; greater amplitudes and/or more frequently periodicity induces stronger alignment (Tuft et al., 2013). To quantify the extent of neurite alignment to the unidirectional patterns, we measured the overall length of the neurite relative to the distance of a straight line drawn parallel with the micropattern from the cell body to the end of the neurite terminus using ImageJ as previously described (Clarke et al., 2011; Tuft et al., 2013). A ratio of neurite length to end-to-end distance with values close to one indicates that the neurite closely follows the pattern whereas a ratio significantly greater than one implies that the neurite deviates from the pattern.

In the first set of experiments, SGNs were cultured on smooth polymer surfaces coated uniformly with peptide, smooth polymer surfaces coated with repeating linear peptide patterns, and polymer surfaces with repeating linear topographical patterns coated uniformly with peptide. On smooth polymer surfaces with no topographical or peptide patterns, SGN neurites wandered randomly evidenced by their high alignment ratio of 3.79±0.28 (Fig. 2A,D). LN and topographical patterning independently improved neurite alignment significantly compared to uniformly coated controls with much lower alignment ratios of 1.47±0.068 (One way ANOVA p<0.01) and 1.45±0.686 (One way ANOVA p<0.01), respectively (Fig 2B,C,D). The alignment ratio was slightly better on topographically patterned substrates compared to substrates with LN patterning, however, this difference was not significant (One way ANOVA p=0.89). It is difficult to make direct comparisons of the relative efficiency of topographical and LN patterning to induce SGN neurite alignment based solely on these data. This is because the level of neurite alignment to topographical patterns varies based on the surface characteristics of amplitude and periodicity (Tuft et al., 2013).

**Figure 2.**
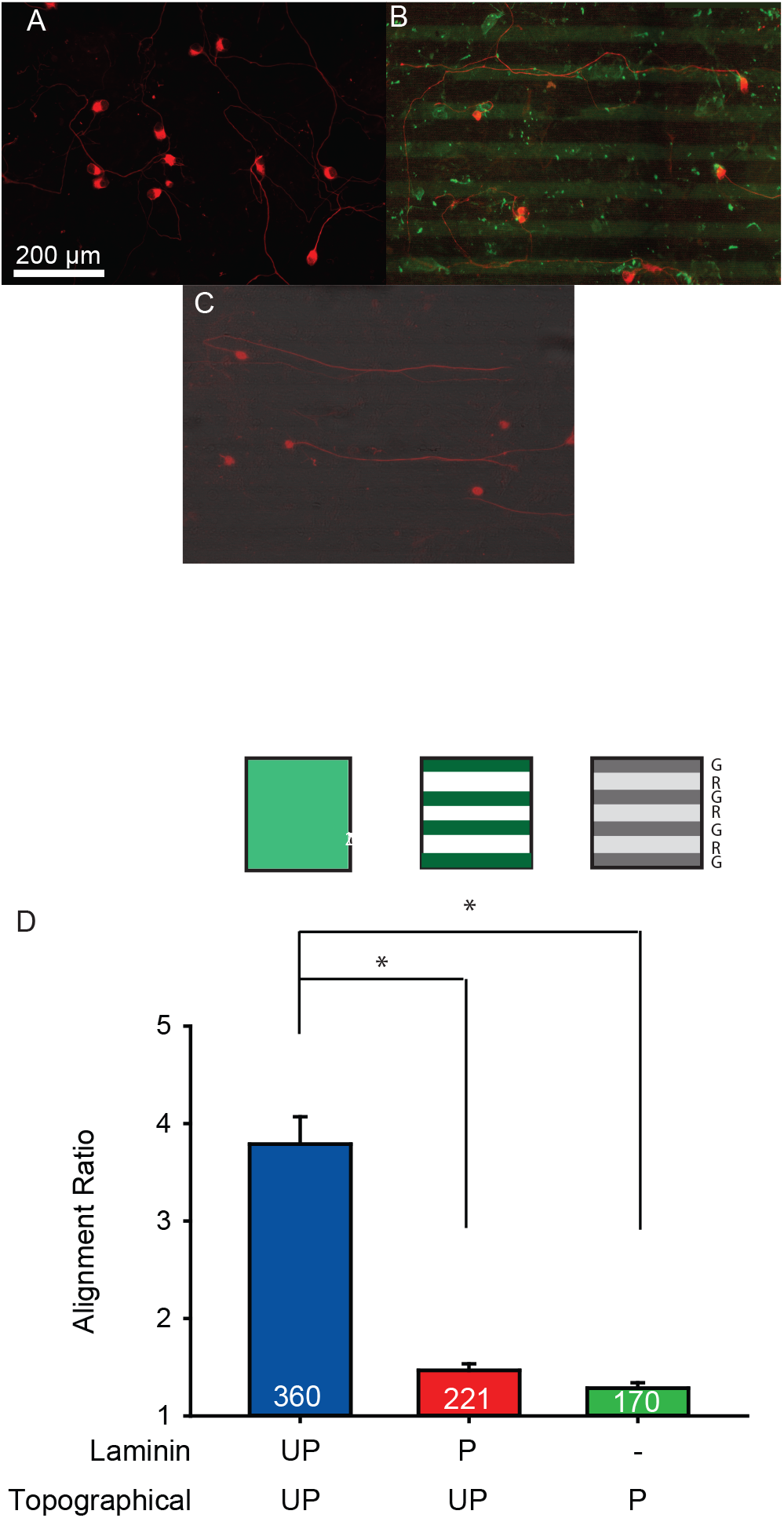
Guidance of dissociated SGN neurite growth on unpatterned (UP) or patterned (P) HMA/HDDMA with or without biochemical patterns of LN. **A-C**. Immunofluorescent images of dissociated SG cultures labeled with anti-NF200 antibody grown on smooth polymers (A), laminin stripes on smooth polymer (B), topographical patterns without LN (C). Neurite growth on unpatterned polymer is random whereas neurites grow parallel to topographically or chemically patterned polymers. Neurites show strong alignment to biochemical LN stripes and stronger alignment to grooves when presented separately. **D**. Mean (±SEM) ratios of neurite length to end-to-end distance for neurites on unpatterned HMA/HDDMA are significantly greater than alignment ratios of both chemical patterning and topographical patterning (One way ANOVA *p<0.01).

### 3.3 SGN neurite alignment can be enhanced or disrupted depending on the site of LN coating

Previous data demonstrate the preference of SGN neurites to grow in the grooves of topographically patterned polymers, implying that encountering the ridges repel the growth cone (S. Li et al., 2016; S. Li et al., 2015). LN was printed on either ridges or grooves of the topographical patterns to assess how biochemical patterns interact with topographical patterns when placed in presumed complimentary (LN in grooves) or antagonistic (LN on ridges) configurations. Specifically, we quantified alignment ratios for neurites grown on topographically patterns polymers with LN patterned in the following ways: uniformly coated (unpatterned), patterned with LN placed in the grooves, or patterned with LN on the ridges. Figure 3 presents alignment ratios for these three conditions. Uniformly coated LN on a topographically patterned polymer yielded an alignment ratio of 1.32±0.09. LN placed in grooves improved neurite alignment with a ratio of 1.08±0.01 whereas LN placed in the ridges reduced the neurite alignment to a ratio of 1.67±0.08.

**Figure 3.**
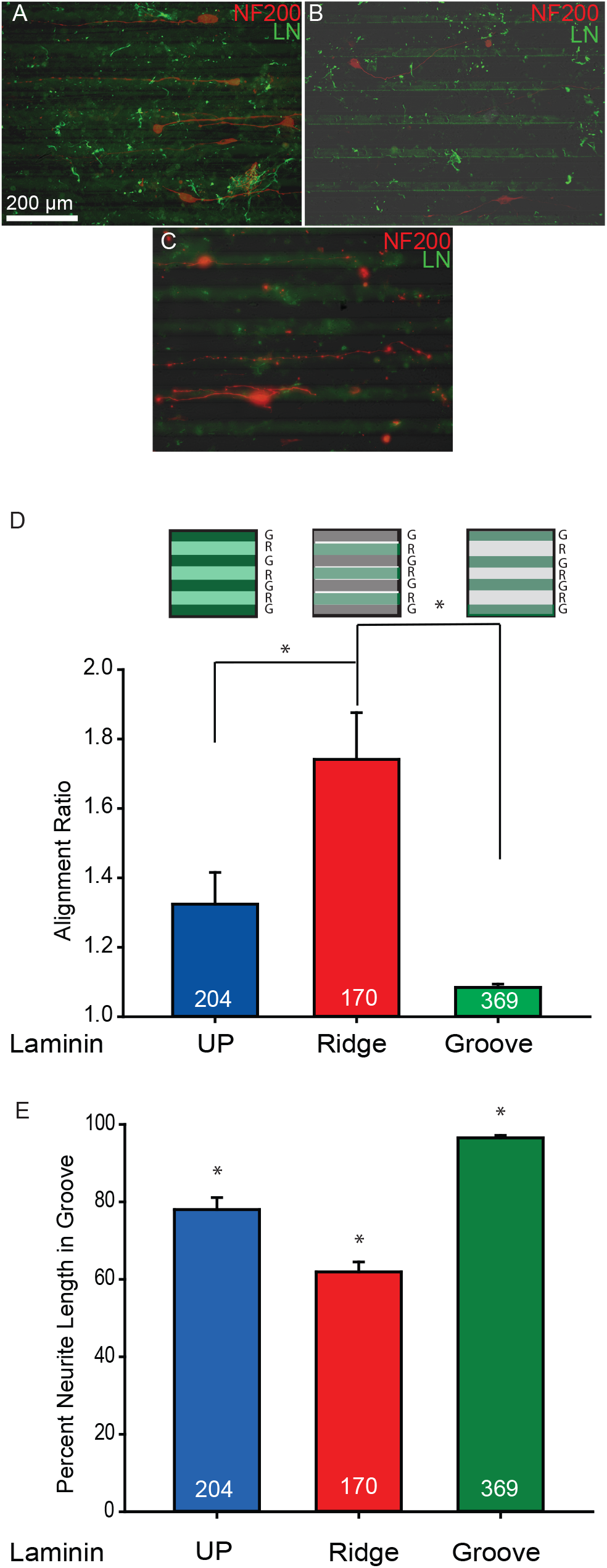
Guidance of dissociated SGN neurite grown on combination topographical and chemical patterns. **A-C**. Immunofluorescent images of dissociated SG cultures grown on topographically patterned surfaces with unpatterned (UP) LN (A), LN on ridges (B) and LN in grooves (C). Neurites show strong alignment to unpatterned LN and stronger alignment to LN patterned in the grooves. Alignment is disrupted when the LN is patterned on the ridges. **D**. Mean (±SEM) ratios of neurite length to end-to-end distance for neurites on patterned HMA/HDDMA with LN patterned on ridges are significantly different than alignment ratios of both LN patterned in grooves and LN unpatterned (One-way ANOVA *p<0.01). **E**. Representation of the percent of neurites in the groove compared to the ridge on topographically patterned surfaces with LN on the ridges, grooves or both. Mean (±SEM) percent neurite lengths in groove are all significantly different (One-way ANOVA *p<0.01).

To further assess the interaction of neurites with combined LN and topographical patterns and how the raised and depressed features of micropatterned substrates and chemical cues influence neurite guidance, the percent of the total neurite length in the groove was also quantified in the presence of different patterning conditions, including uniformly coated LN, LN in the grooves, and LN on the ridges. We measured the length of the neurite when it was in the groove and divided this by the overall length of the neurite. On substrates with uniformly coated LN, neurites showed a strong preference for grooves, with 77.99%±2.01 of the length present in the grooves (Fig. 3). The complimentary addition of LN to the grooves dramatically increased the length of neurite present in the depressed groove features (96.52%±0.59), whereas the antagonistic application of LN to the ridges significantly reduced the length in the grooves (61.89%±2.53) (Fig. 3, One way ANOVA p<0.01). Nevertheless, we see that even in this unfavorable configuration of biochemical patterning, more than 50% of the neurite remains in the groove.

### 3.4 Neural pathfinding on multidirectional micropatterns

Next, we explored the ability of SGN neurites to follow more complex, 90° angle multidirectional topographies. In agreement with prior data, there was a slight skew of neurite segment angles towards lower angles on the repeated 90° angle multidirectional topographical pattern uniformly coated with LN (Fig. 4F) (Tuft et al., 2014). Similarly, on a smooth substrate with LN printed in the multidirectional 90° angle patterns, more than 50% of SGN neurite segments aligned within 10° of the horizontal (Fig. 4E). These data suggest that SGN neurites on either topographical or LN multidirectional 90° angle patterns tend to follow the horizontal and fail to precisely track the pattern angles. Similarly, when LN was printed on ridges of the multidirectional topographical patterned substrates, a majority of the neurite segments aligned to the overall horizontal direction (Fig. 4G). When LN was printed in grooves, a greater percentage of neurite segments with alignment angles closer to 45° were found (Fig. 4H), suggesting that a greater percentage followed the alternating angles of the pattern compared to conditions with uniform LN or LN on the ridges. Significantly, in our previous work, SGNs grown on these multidirectional patterns with uniform LN coating failed to track the 90° angles, even for patterns with higher amplitude features (7.4±0.7 μm) (Tuft et al., 2014). Thus, LN coating in grooves may increase the ability of SGN neurites to follow more complex patterns, whereas simply increasing the amplitude of the topographic features fails to do so. ANOVA results indicate no significant difference in the percentage of neurites between conditions at a given angle, but that there is a significant difference between different angles within a given condition.

**Figure 4.**
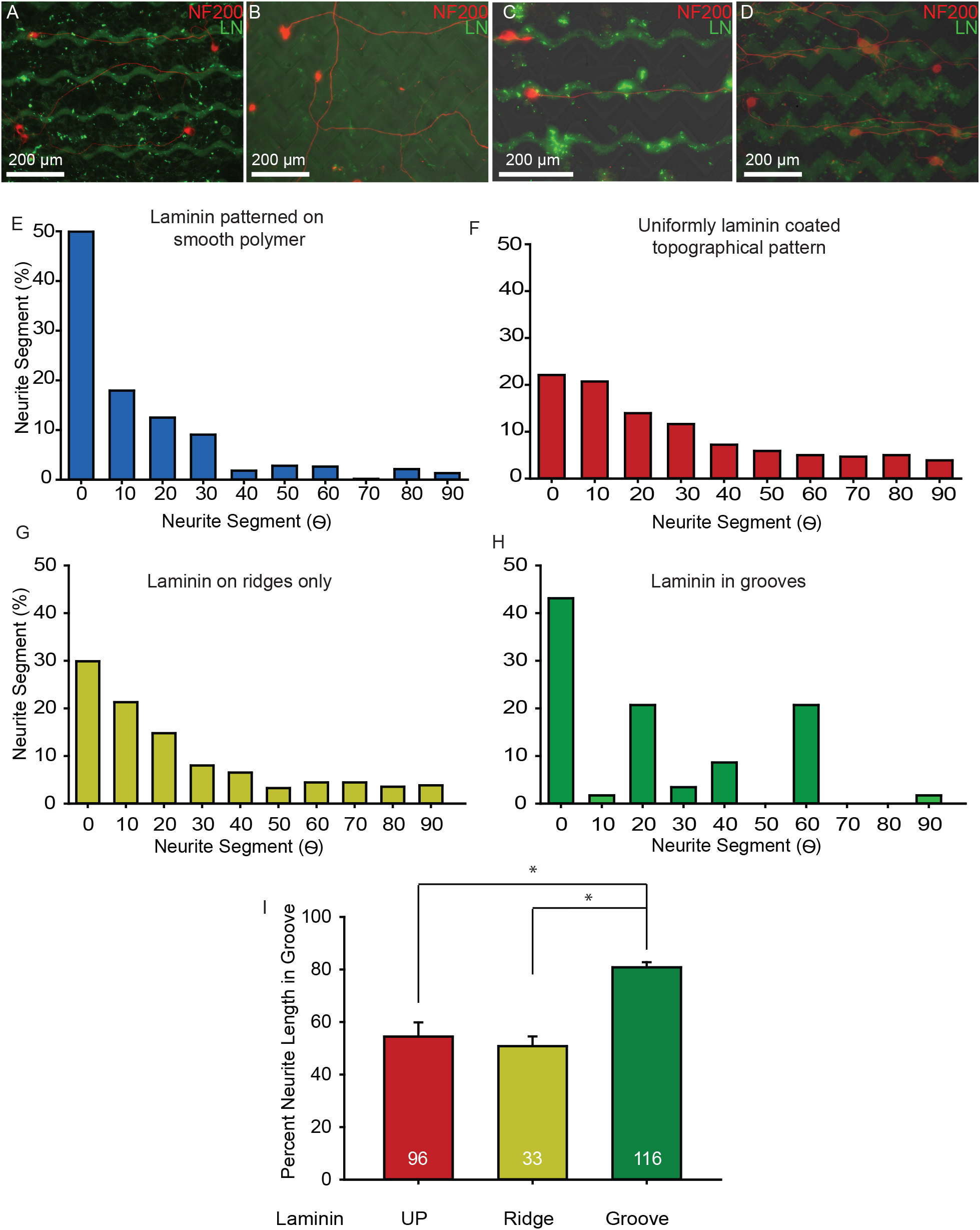
SGN neurite alignment on multidirectional micropatterns. **A-D**. Immunofluorescent images of neurite growth from dissociated SGs on LN patterned on smooth polymers (A), uniformly LN coated on topographically patterned polymers (B), LN printed on either the ridges (C) or grooves (D) of 90° angle substrates. Neurites align to the horizontal in the direction of the pattern without consistent turning at the acute angles. **E-H**. Distribution of SGN neurite segment angles relative to the horizontal plane on LN patterned on smooth polymers (E, n=132), uniformly LN coated on topographically patterned polymers (F, n=110), LN printed on either the ridges (G, n=43) or grooves (H, n=387) of 90° angle substrates. Neurites do not consistently track multidirectional cues as demonstrated by the low incidence of 45° angle neurite segments. They do align somewhat to the horizontal plane with more neurite segments between 0 and 20 degrees. Neurites on LN printed in grooves of 90° angle substrates have the strongest trend to following the horizontal with the neurite segment angles clustered around 0, 20 and 60 degrees. **I**. SGN percent neurite length in depressed microfeatures on multidirectional topographic cues with uniformly LN coated, LN on ridges and LN in grooves. The majority of SGN neurite length is located in grooves in all conditions and a greater neurite length is spent in the grooves when LN is printed in the grooves (One way ANOVA *p<0.01). UP indicates an unpatterned polymer.

Percentage of neurite length in the groove was also determined for neurites on these multidirectional patterns. Topographically patterned substrates with uniformly LN coated showed that neurites remained in the groove ~50% of the time (Fig. 4I). The restriction of LN to ridges did not significantly change the percentage neurite length in grooves compared to controls (One way ANOVA p=0.57) (Fig. 4I). However, the restriction of LN to grooves significantly increased the percentage of neurite length in the grooves to nearly 80% (One way ANOVA p<0.01) (Fig. 4I). These data further demonstrate that complimentary patterning of LN and topographic features (e.g. LN in grooves) enhances the degree of neurite guidance compared to the guidance provided by either patterning cue alone.

### 3.5 Patterning of EphA-4-Fc directs SGN neurite growth and enhances neurite guidance by topographical features

The prior experiments examined the influence of patterning of LN, which supports SGN neurite growth, on SGN guidance. To further explore the ability of patterned biochemical cues to guide SGN neurite growth and to interact with topographical features, we generated micropatterns of EphA4-Fc, which functions to repel SGN neurite growth. In these experiments, stripes of EphA4-Fc with a periodicity of 50 μm were printed on polymers previously coated uniformly with LN. Outgrowth of SGN neurites was random on substrates with uniform LN and uniform EphA4-Fc coating (unpatterned), yielding an alignment ratio of 2.68±0.30 (Fig. 5A,C). On substrates with EphA4-Fc patterning, SGN neurites were almost exclusively found on the LN stripes, between the EphA4-Fc stripes and resulted in an alignment ratio of 1.11±0.02 (p<0.01, Student’s unpaired two-tailed *t*-test) (Fig. 5B, C). Thus, in this system EphA4-Fc appears to provide a chemorepulsive cue capable of precisely directing SGN neurite growth. It is important to note that there is no difference in SGN neurite length when grown on uniform coatings of LN and EphA4-Fc (Sup. Fig. 1).

**Figure 5.**
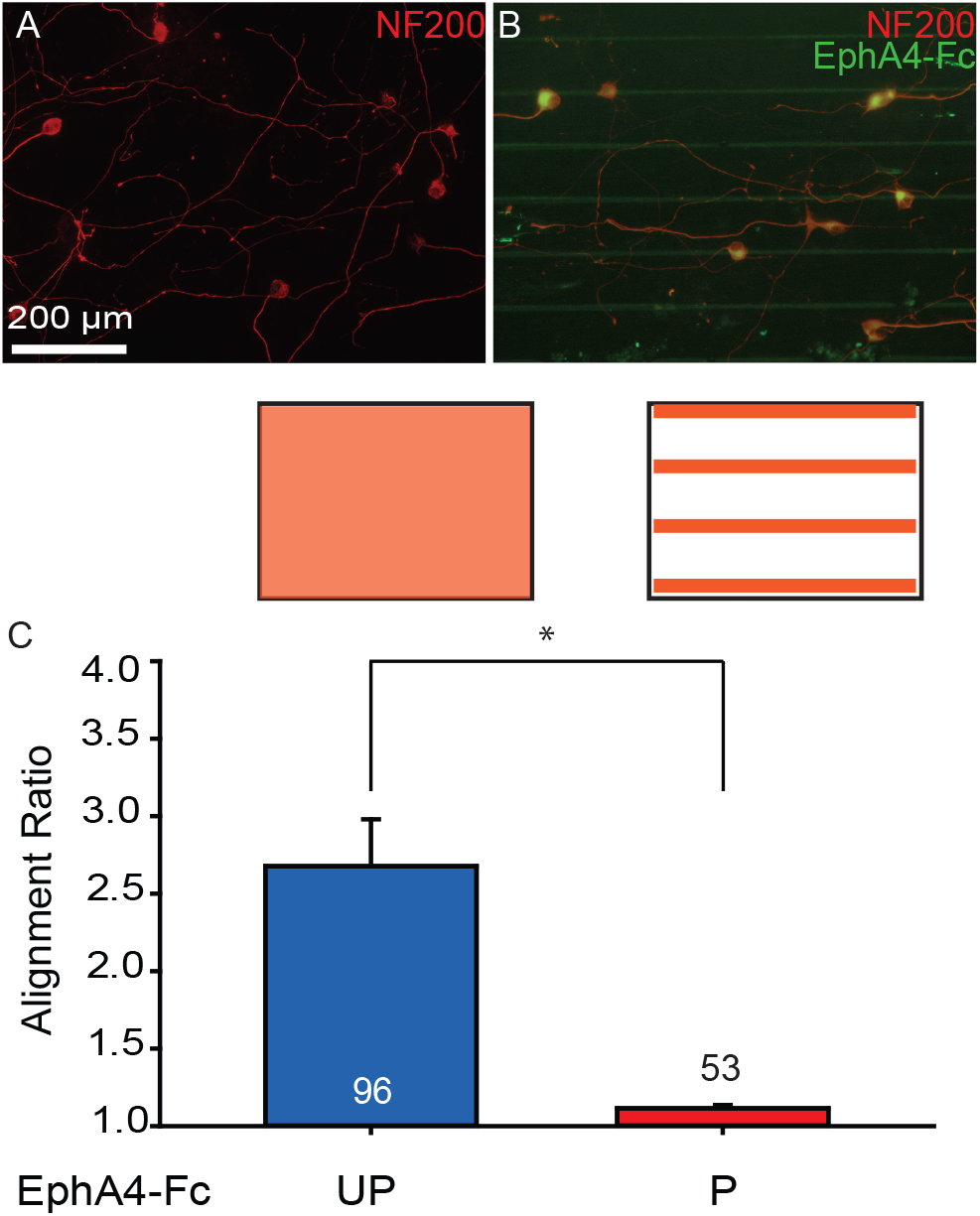
EphA4-Fc and LN show synergistic effects on neurite guidance for topographically unpatterned but biochemically patterned polymers. **A-B**. Immunofluorescent images of neurite growth from dissociated SGNs on uniformly LN and EphA4-Fc coated smooth polymer (A), and uniformly LN coated on smooth polymer with EphA4-Fc stripes (B). Neurite growth on the unpatterned peptides is random whereas neurites on uniformly LN coated on smooth polymer with EphA4-Fc stripes were found between EphA4-Fc stripes. (C) Mean (±SEM) ratios of neurite length to end-to-end distance for neurites on unpatterned HMA/HDDMA are significantly different than alignment ratios of both chemical patterning (*t*-test *p<0.01). P indicates an EphA4-Fc pattern and UP indicates uniformly EphA4-Fc coated substrates.

Next, we examined the influence of LN and EphA4-Fc patterning in combination with topographical features on SGN neurite pathfinding. In this case, three different configurations were created. First, substrates with unidirectional topographical features of 50 μm periodicity and 2 μm amplitude were uniformly coated with EphA4-Fc. Second, LN was selectively placed in the grooves and EphA4-Fc was printed on the ridges of unidirectional topographical patterns. Finally, the opposite configuration was generated with LN on the ridges and EphA4-Fc in the grooves of the unidirectional topographical patterns. SGN neurite growth on topographically patterned substrates coated uniformly with EphA4-Fc resulted in an alignment ratio of 1.44±0.10 (Fig. 6A,D), comparable to the alignment of SGN neurites on the same topographical patterns with a uniform LN coating (1.45±0.686, Fig. 2D). Thus, topographical features guide SGN neurite growth in the presence of different uniform peptide coatings.

**Figure 6.**
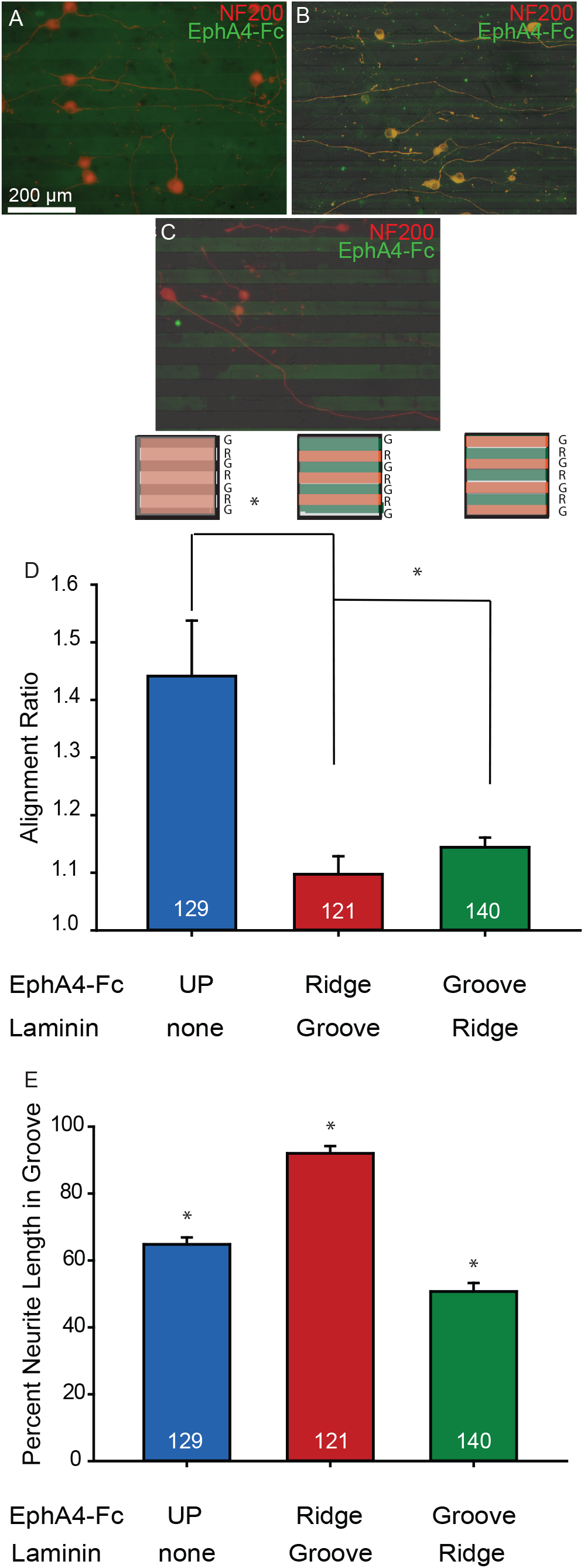
EphA4-Fc improves LN directed SGN neurite guidance on combination topographical and biochemical patterned substrate. **A-C**. Immunofluorescent images of dissociated SG cultures grown on topographically patterned surfaces with uniformly EphA4 coated (A), EphA4 on ridges/LN in grooves (B) and EphA4 in grooves/LN on ridges (C). Neurites show weaker alignment to uniformly EphA4-Fc coated samples and stronger alignment to LN patterned in the ridges and EphA-Fc patterned in grooves. Alignment is strongest when the LN is patterned on the grooves and EphA4-Fc is patterned on the ridges. **D**. Mean (±SEM) ratios of neurite length to end-to-end distance for neurites on patterned HMA/HDDMA with EphA4-Fc patterned on ridges and LN patterned in grooves are significantly different than alignment ratios of both LN patterned on ridges and EphA-Fc patterned in grooves and uniformly EphA4-Fc coated (One way ANOVA *p<0.01). **E**. Representation of the percent of neurites in the groove compared to the ridge on topographically patterned surfaces with EphA4-Fc on the ridges, grooves or both in a background of LN. Mean (±SEM) percent neurite length in groove are all significantly different (One way ANOVA *p<0.01). UP indicates a uniformly EphA4-Fc coated polymer.

Compared to uniform coating of EphA4-Fc, selective placement of LN in the grooves and coating the ridges with EphA4-Fc enhanced SGN neurite alignment with a ratio of 1.09±0.03 (p=0.002); however this was not significantly different than the alignment ratio with LN in the grooves without the EphA4-Fc coating on the ridges (see Fig. 2D, 1.08±0.01, One way ANOVA p=0.59). It is possible that SGN neurites had already reached near perfect alignment on substrates with 2 μm amplitude features and LN coating in the grooves so that coating the ridges with EphA4-Fc did not further enhance neurite alignment in this system. Selective placement of LN on the ridges and coating the grooves with EphA4-Fc disrupted SGN neurite alignment to the pattern (1.14±0.01, One-way ANOVA p=0.003).

To further assess the interaction of neurites with the topographical features and these biochemical cues, the percentage of neurite length in the grooves was also quantified in the presence of these different biochemical patterning conditions. In the presence of uniform EphA4-Fc, neurites showed a preference for grooves, with 64.78%±2.09 of the length being present in the grooves (Fig. 6E). The addition of LN to the grooves and EphA4-Fc to the ridges increased the length of neurite present in the groove (91.98%±1.95), whereas application of LN to the ridges and EphA4-Fc to the grooves reduces the length of neurite in the groove (50.74%±2.51). These results are consistent with the notion that the raised topographical features function as repulsive cues leading to a preference for neurites to remain in the grooves (S. Li et al., 2015). Coating the grooves with EphA4-Fc, which repels neurite growth, and the ridges with LN, which facilitates neurite growth, presents the growth cone with conflicting topographical and biochemical guidance cues and disrupts neurite alignment to some degree.

## 4. Discussion

Photopolymerization and microcontact printing/selective absorption of peptides can be used to generate biochemical and topographical features in defined patterns that enhance or disrupt SGN neurite alignment to the features (Fig 1). Observations from these studies can be used to infer the hierarchical relationships of the biochemical and topographical guidance cues in this system. First, these results demonstrate that SGN neurites preferentially grow on LN-coated stripes and avoid EphA4-Fc coated stripes so that unidirectional patterns of either peptide direct SGN neurite pathfinding (Fig. 2 and Fig. 5). Second, our data demonstrate that a combination biochemical and topographical patterning in parallel can enhance or disrupt neurite guidance depending on where the peptides are placed (Fig. 3 and Fig. 6). Specifically, we found that coating the ridges with EphA4-Fc provided further repulsion of neurite growth across the feature and lead to enhanced alignment (Fig 6). Likewise, coating the grooves with LN enhanced neurite alignment to both unidirectional and multidirectional topographies (Fig 3, Fig 4, and Fig 6). Conversely, we found reduced neurite alignment on micropatterns with EphA4-Fc coating in the grooves or LN coating on the ridges correlated with a significantly reduced preference for the neurite to grow in the grooved aspect of the topographically features (Fig. 6). These data further support the notion that the complimentary application of biochemical patterns to topographical patterns enhances neurite alignment, as is the case when LN was added selectively to grooved features or EphA4-Fc to the ridges. Conversely, the application of antagonistic biochemical patterns disrupts neurite alignment to the micropatterns. Apparently the ‘repulsive’ ridges become more favorable for neurite growth if coated with a biochemical cue that supports neurite growth and this thereby facilitates neurites to cross ridge features and decrease alignment to the micropatterns.

These data can also be used to infer suggesting the hierarchical dominance of the topographical cues over the LN cues with these particular topographical feature dimensions. For example, the majority of SGN neurites remained in the grooves when the ridges are coated with LN (Fig. 3E). This suggests that topographical features of (2 μm amplitude and 50 μm periodicity) exert a greater influence on neurite growth than the LN coating. Likewise, SGN neurites tend to remain in the grooves even when the groove is coated with chemorepulsive EphA4-Fc peptide implying that the topographical features in this study dominate over this chemorepulsive cue (Fig. 6E). It is important to note that these observations apply to the specific topographical features and surface biochemical coatings examined here (2μm amplitude and 50μm periodicity), and the relative dominance of topographical or biochemical patterns may change as feature dimensions and/or biochemical surface coating are altered.

Lastly, we have previously reported EphA4-Fc appears to function as a chemorepulsive signal when the SGN neurites face a transition from a different substrate (e.g. LN) to EphA4-Fc (Tuft et al., 2018). Interestingly in this study we show that SGN neurites grow well on substrates uniformly coated with EphA4-Fc and that without this transition or border, SGN neurites can grow on EphA4-Fc normally (Fig. 5 and Sup. Fig. 1). The chemorepulsive nature of EphA4-Fc only arises when neurites face a transition from LN to EphA4-Fc in our experience.

The long-term goal of these studies is to achieve spatially organized regeneration of SGN peripheral processes to significantly reduce the physical distance between SGNs and CI electrodes (Senn et al., 2017). A regenerating neurite faces a complex milieu of physical, cellular, and biochemical cues that interact to determine the ultimate trajectory of neurite growth. These studies focus on the interplay between specific topographical features and biochemical cues that have previously been demonstrated in isolation to guide neurite growth. The results inform the ultimate the goal of engineering scaffolds with topographical and/or biochemical features sufficient to support and precisely guide growth of regenerating SGN fibers. For this to occur, neurite regeneration must first be stimulated. Several groups have demonstrated that neurotrophic factors, such as BDNF and NT-3, are sufficient to induce SGN neurite regeneration *in vivo* following deafening (Budenz et al., 2015; H. Li et al., 2017; Shibata et al., 2010; Wise et al. 2005). Further, gradients of neurotrophic factors have been shown to guide SGN neurite growth in vitro (H. Li et al., 2017). Once neurite regeneration has been stimulated, scaffoldings engineered to guide neurite growth could then be deployed to ensure that the neurite growth remains organized and is directed towards intended targets (Ceschi et al., 2014; H. Li et al., 2017). Another critical step will be to provide “stop” signals that arrest growth near the desired target (Roehm & Hansen, 2005). Finally, for the regenerated fibers to fire properly and remain healthy it will likely prove critical to ensure that they become myelinated with Schwann cells, thereby reflecting their status in the healthy cochlea.

SGN are bipolar neurons with a peripheral axon that projects to the organ of Corti to innervate hair cells and a central process that projects to the cochlear nucleus. While the focus of this work is to understand how physical and biochemical cues interact to direct the regeneration of SGN peripheral processes (e.g. towards a stimulating electrode), the results reported here are derived from dissociated SGN cultures and are not able to distinguish putative central from peripheral processes. As molecular markers become available to differentiate central from peripheral processes it will be important to determine the extent to which the neurites from these dissociated cultures reflect properties of peripheral SGN axons *in vivo*. Prior work with explant cultures, in which it is possible to distinguish peripheral and central process based on the morphology of the explant, suggests that SGN peripheral processes respond to the types of topographical and chemical cues used in these experiments (Brors et al., 2003; Clarke et al., 2011).

Another limitation of the current study is that the neurons used were derived from neonatal animals yet the target neurons for CI electrodes are mature. Importantly, recent work indicates that neurites from adult SGNs and adult dorsal root ganglion neurons respond to similar topographic features as those used in these studies (L. Xu et al., 2018) lending confidence that such features will prove effective in mature cochleae.

## 5. Conclusions

This paper presents methods to generate substrates with independently fabricated surface topographical and biochemical patterns. When presented with either the topographical or the biochemical patterns alone, SGN neurites significantly aligned to the pattern presented compared to unpatented controls. Combination of biochemical cues that act in cooperation with topographical features enhances neurite alignment to the features whereas combination of biochemical cues that present antagonist cues to the topographical features disrupts neurite alignment. Thus, systems of combined biochemical and topographical patterning can likely be engineered to precisely direct regeneration of target nerve fibers towards the stimulating electrodes in an effort to enhance the spatial resolution provided by neural prostheses.

## Abbreviations

LN: laminin
MA: methacrylate
HMA: hexyl methacrylate
HDDMA: hexanediol dimethacrylate

---

**Supplemental Figure 1.** SGN neurite length is equal unaffected by peptide substrate. Mean neurite length of SGNs grown on topographically patterned substrates with uniform coatings of either LN of EphA4-Fc. No significant difference was found in neurite length between substrate. (t-test p = 0.55)

## References

Aletsee, C., Brors, D., Palacios, S., Pak, K., Mullen, L., Dazert, S., & Ryan, A. F. (2002). The effects of laminin-1 on spiral ganglion neurons are dependent on the MEK/ERK signaling pathway and are partially independent of Ras. Hearing research, 164(1–2), 1–11.

Apple, D. J., & Sims, J. (1996). Harold Ridley and the invention of the intraocular lens. Surv Ophthalmol, 40(4), 279–292.

Belisle, J. M., Correia, J. P., Wiseman, P. W., Kennedy, T. E., & Costantino, S. (2008). Patterning protein concentration using laser-assisted adsorption by photobleaching, LAPAP. Lab Chip, 8(12), 2164–2167. doi:10.1039/b813897d

Branch, D. W., Corey, J. M., Weyhenmeyer, J. A., Brewer, G. J., & Wheeler, B. C. (1998). Microstamp patterns of biomolecules for high-resolution neuronal networks. Med Biol Eng Comput, 36(1), 135–141.

Branch, D. W., Wheeler, B. C., Brewer, G. J., & Leckband, D. E. (2001). Long-term stability of grafted polyethylene glycol surfaces for use with microstamped substrates in neuronal cell culture. Biomaterials, 22(10), 1035–1047.

Brors, D., Bodmer, D., Pak, K., Aletsee, C., Schafers, M., Dazert, S., & Ryan, A. F. (2003). EphA4 provides repulsive signals to developing cochlear ganglion neurites mediated through ephrin-B2 and-B3. Journal of Comparative Neurology, 462(1), 90–100. doi:10.1002/cne.10707

Budenz, C. L., Wong, H. T., Swiderski, D. L., Shibata, S. B., Pfingst, B. E., & Raphael, Y. (2015). Differential effects of AAV.BDNF and AAV.Ntf3 in the deafened adult guinea pig ear. Sci Rep, 5, 8619. doi:10.1038/srep08619

Ceschi, P., Bohl, A., Sternberg, K., Neumeister, A., Senz, V., Schmitz, K. P., … Paasche, G. (2014). Biodegradable polymeric coatings on cochlear implant surfaces and their influence on spiral ganglion cell survival. J Biomed Mater Res B Appl Biomater, 102(6), 1255–1267. doi:10.1002/jbm.b.33110

Clarke, J. C., Tuft, B. W., Clinger, J. D., Levine, R., Figueroa, L. S., Allan Guymon, C., & Hansen, M. R. (2011). Micropatterned methacrylate polymers direct spiral ganglion neurite and Schwann cell growth. Hearing research, 278(1–2), 96–105. doi:10.1016/j.heares.2011.05.004

Clarke, J. C., Tuft, B. W., Clinger, J. D., Levine, R., Figueroa, L. S., Guymon, C. A., & Hansen, M. R. (2011). Micropatterned methacrylate polymers direct spiral ganglion neurite and Schwann cell growth. Hear Res, 278(1–2), 96–105. doi:10.1016/j.heares.2011.05.004

Coate, T. M., Raft, S., Zhao, X., Ryan, A. K., Crenshaw, E. B., 3rd, & Kelley, M. W. (2012). Otic mesenchyme cells regulate spiral ganglion axon fasciculation through a Pou3f4/EphA4 signaling pathway. Neuron, 73(1), 49–63. doi:10.1016/j.neuron.2011.10.029

Defourny, J., Poirrier, A. L., Lallemend, F., Mateo Sanchez, S., Neef, J., Vanderhaeghen, P., … Malgrange, B. (2013). Ephrin-A5/EphA4 signalling controls specific afferent targeting to cochlear hair cells. Nature Communications, 4, 1438. doi:10.1038/ncomms2445

Evans, A. R., Euteneuer, S., Chavez, E., Mullen, L. M., Hui, E. E., Bhatia, S. N., & Ryan, A. F. (2007). Laminin and fibronectin modulate inner ear spiral ganglion neurite outgrowth in an in vitro alternate choice assay. Dev Neurobiol.

Fricke, R., Zentis, P. D., Rajappa, L. T., Hofmann, B., Banzet, M., Offenhausser, A., & Meffert, S. H. (2011). Axon guidance of rat cortical neurons by microcontact printed gradients. Biomaterials, 32(8), 2070–2076. doi:10.1016/j.biomaterials.2010.11.036

Gustavsson, P., Johansson, F., Kanje, M., Wallman, L., & Linsmeier, C. E. (2007). Neurite guidance on protein micropatterns generated by a piezoelectric microdispenser. Biomaterials, 28(6), 1141–1151.

Hansen, M. R., Vijapurkar, U., Koland, J. G., & Green, S. H. (2001). Reciprocal signaling between spiral ganglion neurons and Schwann cells involves neuregulin and neurotrophins. Hearing research, 161(1–2), 87–98.

Hansen, M. R., Zha, X. M., Bok, J., & Green, S. H. (2001). Multiple distinct signal pathways, including an autocrine neurotrophic mechanism, contribute to the survival-promoting effect of depolarization on spiral ganglion neurons in vitro. The Journal of neuroscience: the official journal of the Society for Neuroscience, 21(7), 2256–2267.

Honegger, T., Thielen, M. I., Feizi, S., Sanjana, N. E., & Voldman, J. (2016). Microfluidic neurite guidance to study structure-function relationships in topologically-complex population-based neural networks. Sci Rep, 6, 28384. doi:10.1038/srep28384

Jeon, E. J., Xu, N., Xu, L., & Hansen, M. R. (2011). Influence of central glia on spiral ganglion neuron neurite growth. Neuroscience, 177, 321–334. doi:10.1016/j.neuroscience.2011.01.014

Johansson, F., Carlberg, P., Danielsen, N., Montelius, L., & Kanje, M. (2006). Axonal outgrowth on nano-imprinted patterns. Biomaterials, 27(8), 1251–1258.

Joo, S., Kang, K., & Nam, Y. (2015). In vitro neurite guidance effects induced by polylysine pinstripe micropatterns with polylysine background. J Biomed Mater Res A, 103(8), 2731–2739. doi:10.1002/jbm.a.35405

Kenny, S. M., & Buggy, M. (2003). Bone cements and fillers: a review. J Mater Sci Mater Med, 14(11), 923–938.

Kijenska-Gawronska, E., Bolek, T., Bil, M., & Swieszkowski, W. (2019). Alignment and bioactive molecule enrichment of bio-composite scaffolds towards peripheral nerve tissue engineering. Journal of Materials Chemistry B, 7(29), 4509–4519. doi:10.1039/c9tb00367c

Klein, C. L., Scholl, M., & Maelicke, A. (1999). Neuronal networks in vitro: formation and organization on biofunctionalized surfaces. J Mater Sci Mater Med, 10(12), 721–727.

Leach, J. B., Achyuta, A. K., & Murthy, S. K. (2010). Bridging the Divide between Neuroprosthetic Design, Tissue Engineering and Neurobiology. Frontiers in neuroengineering, 2, 18. doi:10.3389/neuro.16.018.2009

Li, H., Edin, F., Hayashi, H., Gudjonsson, O., Danckwardt-Lilliestrom, N., Engqvist, H., … Xia, W. (2017). Guided growth of auditory neurons: Bioactive particles towards gapless neural - electrode interface. Biomaterials, 122, 1–9. doi:10.1016/j.biomaterials.2016.12.020

Li, S., Tuft, B., Xu, L., Polacco, M., Clarke, J. C., Guymon, C. A., & Hansen, M. R. (2016). Intracellular calcium and cyclic nucleotide levels modulate neurite guidance by microtopographical substrate features. J Biomed Mater Res A, 104(8), 2037–2048. doi:10.1002/jbm.a.35738

Li, S., Tuft, B. W., Xu, L., Polacco, M. A., Clarke, J. C., Guymon, C. A., & Hansen, M. R. (2015). Microtopographical features generated by photopolymerization recruit RhoA/ROCK through TRPV1 to direct cell and neurite growth. Biomaterials, 53, 95–106. doi:10.1016/j.biomaterials.2015.02.057

Li, W., Tang, Q. Y., Jadhav, A. D., Narang, A., Qian, W. X., Shi, P., & Pang, S. W. (2015). Large-scale topographical screen for investigation of physical neural-guidance cues. Sci Rep, 5, 8644. doi:10.1038/srep08644

Millet, L. J., Stewart, M. E., Nuzzo, R. G., & Gillette, M. U. (2010). Guiding neuron development with planar surface gradients of substrate cues deposited using microfluidic devices. Lab on a Chip, 10(12), 1525–1535. doi:10.1039/c001552k

Nadol, J. B., Jr., Shiao, J. Y., Burgess, B. J., Ketten, D. R., Eddington, D. K., Gantz, B. J., … Shallop, J. K. (2001). Histopathology of cochlear implants in humans. Ann Otol Rhinol Laryngol, 110(9), 883–891.

O’Leary, S. J., Richardson, R. R., & McDermott, H. J. (2009). Principles of design and biological approaches for improving the selectivity of cochlear implant electrodes. J Neural Eng, 6(5), 055002. doi:10.1088/1741-2560/6/5/055002

Pettingill, L. N., Richardson, R. T., Wise, A. K., O’Leary, S. J., & Shepherd, R. K. (2007). Neurotrophic factors and neural prostheses: potential clinical applications based upon findings in the auditory system. IEEE Trans Biomed Eng, 54(6 Pt 1), 1138–1148.

Pfingst, B. E., Bowling, S. A., Colesa, D. J., Garadat, S. N., Raphael, Y., Shibata, S. B., … Zhou, N. (2011). Cochlear infrastructure for electrical hearing. Hearing research. doi:10.1016/j.heares.2011.05.002

Roehm, P. C., & Hansen, M. R. (2005). Strategies to preserve or regenerate spiral ganglion neurons. Curr Opin Otolaryngol Head Neck Surg, 13(5), 294–300.

Rubinstein, J. T. (2004). How cochlear implants encode speech. Curr Opin Otolaryngol Head Neck Surg, 12(5), 444–448.

Senn, P., Roccio, M., Hahnewald, S., Frick, C., Kwiatkowska, M., Ishikawa, M., … Lowenheim, H. (2017). NANOCI-Nanotechnology Based Cochlear Implant With Gapless Interface to Auditory Neurons. Otol Neurotol, 38(8), e224–e231. doi:10.1097/MAO.0000000000001439

Shannon, R. V., Fu, Q. J., & Galvin, J., 3rd. (2004). The number of spectral channels required for speech recognition depends on the difficulty of the listening situation. Acta Otolaryngol Suppl(552), 50–54.

Shibata, S. B., Cortez, S. R., Beyer, L. A., Wiler, J. A., Di Polo, A., Pfingst, B. E., & Raphael, Y. (2010). Transgenic BDNF induces nerve fiber regrowth into the auditory epithelium in deaf cochleae. Exp Neurol, 223(2), 464–472. doi:10.1016/j.expneurol.2010.01.011

Tuft, B. W., Li, S., Xu, L., Clarke, J. C., White, S. P., Guymon, B. A., … Guymon, C. A. (2013). Photopolymerized microfeatures for directed spiral ganglion neurite and Schwann cell growth. Biomaterials, 34(1), 42–54. doi:10.1016/j.biomaterials.2012.09.053

Tuft, B. W., Xu, L., Leigh, B., Lee, D., Guymon, C. A., & Hansen, M. R. (2018). Photopolymerized micropatterns with high feature frequencies overcome chemorepulsive borders to direct neurite growth. J Tissue Eng Regen Med, 12(3), e1392–e1403. doi:10.1002/term.2527

Tuft, B. W., Xu, L., White, S. P., Seline, A. E., Erwood, A. M., Hansen, M. R., & Guymon, C. A. (2014). Neural pathfinding on uni- and multidirectional photopolymerized micropatterns. ACS Appl Mater Interfaces, 6(14), 11265–11276. doi:10.1021/am501622a

Uludag, H., De Vos, P., & Tresco, P. A. (2000). Technology of mammalian cell encapsulation. Adv Drug Deliv Rev, 42(1–2), 29–64.

Wise, A. K., Richardson, R., Hardman, J., Clark, G., & O’Leary, S. (2005). Resprouting and survival of guinea pig cochlear neurons in response to the administration of the neurotrophins brain-derived neurotrophic factor and neurotrophin-3. Journal of Comparative Neurology, 487(2), 147–165. doi:10.1002/cne.20563

Wittig, J. H., Jr., Ryan, A. F., & Asbeck, P. M. (2005). A reusable microfluidic plate with alternate-choice architecture for assessing growth preference in tissue culture. J Neurosci Methods, 144(1), 79–89.

Xu, L., Seline, A. E., Leigh, B., Ramirez, M., Guymon, C. A., & Hansen, M. R. (2018). Photopolymerized Microfeatures Guide Adult Spiral Ganglion and Dorsal Root Ganglion Neurite Growth. Otol Neurotol, 39(1), 119–126. doi:10.1097/MAO.0000000000001622

Xu, N., Engbers, J., Khaja, S., Xu, L., Clark, J. J., & Hansen, M. R. (2012). Influence of cAMP and protein kinase A on neurite length from spiral ganglion neurons. Hearing research, 283(1–2), 33–44. doi:10.1016/j.heares.2011.11.010

