## Supplementary figures and images for "Interaction of micropatterned topographical and biochemical cues to direct neurite growth from spiral ganglion neurons"

### Supplemental Figure 1

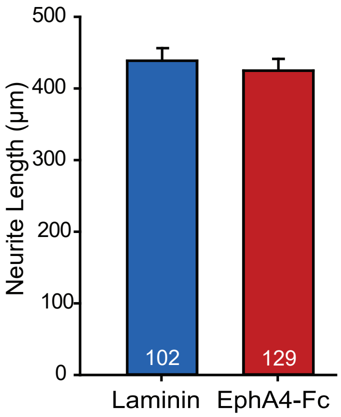
